# Temporal evolution from retinal image size to perceived size in human visual cortex

**DOI:** 10.1101/455139

**Authors:** Juan Chen, Irene Sperandio, Molly J. Henry, Melvyn A Goodale

## Abstract

Our visual system affords a distance-invariant percept of object size by integrating retinal image size with viewing distance (size constancy). Single-unit studies with animals have shown that real changes in distance can modulate the firing rate of neurons in primary visual cortex and even subcortical structures, which raises an intriguing possibility that the required integration for size constancy may occur in the initial visual processing in V1 or even earlier. In humans, however, EEG and brain imaging studies have typically manipulated the apparent (not real) distance of stimuli using pictorial illusions, in which the cues to distance are sparse and not congruent. Here, we physically moved the monitor to different distances from the observer, a more ecologically valid paradigm that emulates what happens in everyday life. Using this paradigm in combination with electroencephalography (EEG), we were able for the first time to examine how the computation of size constancy unfolds in real time under real-world viewing conditions. We showed that even when all distance cues were available and congruent, size constancy took about 150 ms to emerge in the activity of visual cortex. The 150-ms interval exceeds the time required for the visual signals to reach V1, but is consistent with the time typically associated with later processing within V1 or recurrent processing from higher-level visual areas. Therefore, this finding provides unequivocal evidence that size constancy does not occur during the initial signal processing in V1 or earlier, but requires subsequent processing, just like any other feature binding mechanisms.

## Main text

Our visual perception of the world is not a simple reflection of incoming retinal inputs, but involves complex integration of spatial and/or temporal contextual information. One clear example of this integration is size constancy, in which we tend to perceive the size of an object at different distances as constant, even though the image it subtends on the retina (retinal image size) changes with viewing distance. Size constancy requires that we integrate retinal image size with information about viewing distance. When (and where) the computations underlying size constancy take place in the visual brain is an important question as it speaks to when (and where) our brain can infer the physical property of objects in the outside world based on sensory input. A long history of neuropsychological studies has shown that lesions in occipitotemporal, inferotemporal, and parietal cortices interfere with size constancy judgements (Humphrey & Weiskrantz, 1969; Ungerleider, Ganz, & Pribram, 1977; Wyke, 1960). Yet, single-cell recording studies suggest that the required neural components for the computation of size constancy could be present as early as in the primary visual cortex V1 (Dobbins, Jeo, Fiser, & Allman, 1998; Marg & Adams, 1970; Ni, Murray, & Horwitz, 2014; Smith & Marg, 1975) – even though traditionally this visual area has been thought to faithfully code the retinal input. Along these lines, a growing body of evidence from human fMRI studies showed that V1 actually represents perceived size (Fang, Boyaci, Kersten, & Murray, 2008; He, Mo, Wang, & Fang, 2015; Murray, Boyaci, & Kersten, 2006; Pooresmaeili, Arrighi, Biagi, & Morrone, 2013; Sperandio, Chouinard, & Goodale, 2012), which requires the integration of retinal image size and viewing distance.

Although V1 can represent perceived size, there are two possibilities with respect to when (and where) the integration itself might occur. On the one hand, it is possible that the integration of retinal signals and viewing distance (as signaled by a range of possible visual and oculomotor cues) would begin in the initial stages of signal processing in V1 (Pooresmaeili et al., 2013) or even earlier in the thalamus before any signals have even reached the cortex (Richards, 1968; Richards, 1971). Such an early integration is an intriguing possibility because a rapid computation of the real-world size of objects is not only critical for stabilized perception, but is also important for accurate motor planning, where size and distance information have to be integrated for the execution of skilled actions, such as reaching and grasping. In fact, previous neurophysiological studies have shown that ocular vergence, accommodation, gaze direction, and viewing distance (i.e., eye position) could influence the spiking rate of neurons in the lateral geniculate nucleus (LGN) (Lal & Friedlander, 1990; Weyand & Malpeli, 1993) and/or V1 (Dobbins et al., 1998; Marg & Adams, 1970; Masson, Busettini, & Miles, 1997; Rosenbluth & Allman, 2002; Trotter & Celebrini, 1999; Trotter, Celebrini, Stricanne, Thorpe, & Imbert, 1992; Weyand & Malpeli, 1993), and some of the influence could be present at the beginning of the visual response in V1 (Trotter & Celebrini, 1999). On the other hand, it is also possible that the initial signals in V1 represent purely retinal input and any modulation of those signals by viewing distance occurs later (Blakemore, Garner, & Sweet, 1972; Fang et al., 2008; Humphrey & Weiskrantz, 1969; Liu et al., 2009; Ni et al., 2014; Sperandio et al., 2012). Due to the low temporal resolution of blood-oxygen-level dependent (BOLD) signals, functional imaging studies cannot directly address which of these possibilities discussed above is the most likely.

Here, by using high-temporal resolution EEG together with a multivariate pattern analysis of the signals as they unfolded, we focused on the time course of any integration that reflected the operation of size constancy. Recent electrophysiological studies in monkeys (Ni et al., 2014) and humans (Liu et al., 2009) have investigated the timing of the modulation of the representation of size by perceived distance. However, these studies did not systematically explore the effects of changing the *real* distance of the stimulus, but manipulated apparent distance by using the Ponzo illusion instead. In such illusion, the pictorial cues signal that objects are located at different distances from the observer but the binocular and oculomotor cues (vergence and accommodation) always signal a fixed distance, i.e., the real distance of the monitor on which the pictorial illusion is displayed. This incongruence in the distance cues could potentially delay and even interfere with the integration of distance and retinal image size (Sperandio et al., 2012). Thus, to investigate the time course of size-distance integration using EEG, we devised a natural viewing paradigm in which the cues to distance were entirely congruent. To this end, we physically moved the entire visual display to different distances from the observer. Thus, compared to the pictorial illusions that have been used in previous studies (Fang et al., 2008; He et al., 2015; Liu et al., 2009; Murray et al., 2006; Ni et al., 2014; Schwarzkopf, Song, & Rees, 2011), our experiments were much more ecologically valid because they emulated what happens in everyday life when people look at objects located at different distances. We measured EEG activity when size constancy was essentially perfect (Experiments 1 and 2) and when it was disrupted by removing most of the cues to distance (Experiment 3).

## Results

To investigate the temporal evolution of the representation of stimulus size (i.e., retinal image size versus perceived size), the physical size and viewing distance of the visual stimulus were manipulated to create four conditions: near-small (NS), near-large (NL), far-small (FS), and far-large (FL) (**Fig. 1A**). Crucially, the stimuli in the NS and FL conditions had the same retinal size, while those in the NS and FS conditions had the same physical size, as did those in the NL and FL conditions. These relationships between the different conditions in retinal image size and in physical size are reflected in the two “similarity matrices”, shown in **Fig. 1B,** which by definition were the same for all participants. Unlike retinal size or physical size, however, the perceived size of each stimulus depends on the availability and weighting of distance cues (Chen, Sperandio, & Goodale, 2018; Holway & Boring, 1941; Sperandio & Chouinard, 2015) and could vary between individuals [see **Fig. 1B, right column** for an example of the “similarity” in perceived size from one participant in Experiment 3 in which distance cues were restricted (**Fig. 2B left**)].

**Fig 1.**
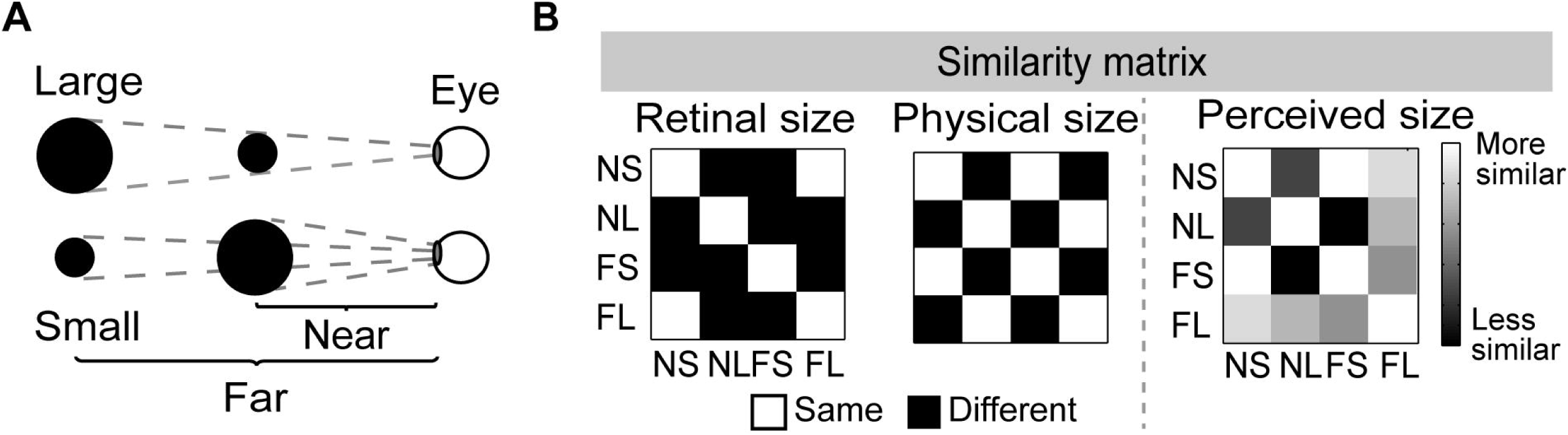
Design of all experiments and the “similarity” matrix between conditions. **A**, Solid circles of two sizes (Small = 4 cm and Large = 8 cm) were presented at two distances (Near = 28.5 cm and Far = 57 cm). **B**, The retinal-image size similarity matrix, the physical-size similarity matrix, and the perceived size similarity matrix for all conditions. The retinal-size and physical-size matrices consisted of values of “0” s (i.e. 0s indicate “different” retinal size or physical size) or “1”s (1s indicate the “same” retinal size or the same physical size). The elements of the perceived size similarity matrix were calculated for each participant based on the “similarity” of the reported perceived size between each pair of conditions. The “similarity” was operationally defined as the difference in perceived size between each pair of conditions multiplied by −1.

**Fig 2.**
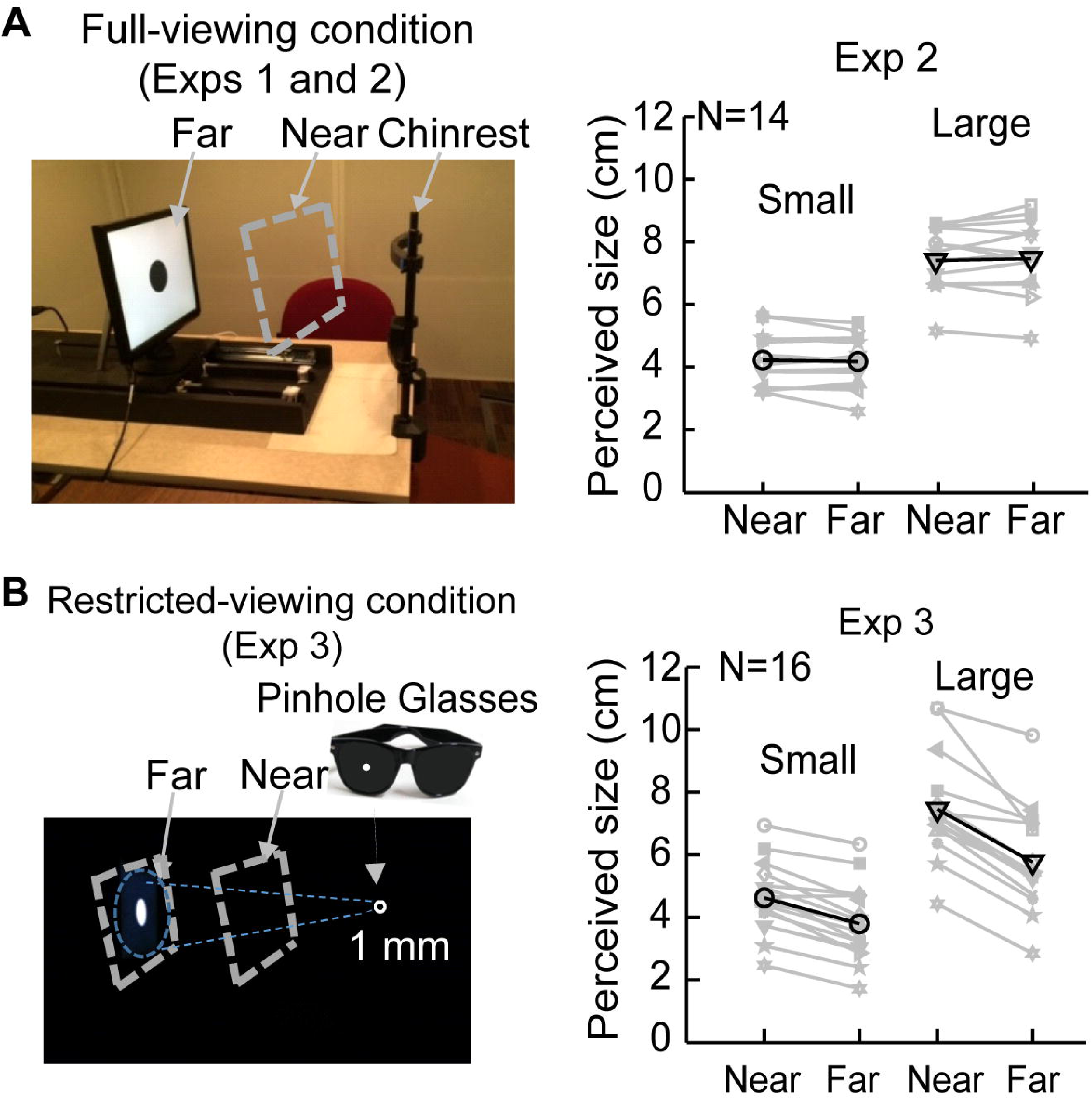
Viewing conditions and the behavioral results of perceived size in the corresponding viewing conditions. **A**, Left: In both Experiments 1 and 2, participants viewed the stimuli binocularly with room lights on (i.e., full-viewing condition). The stimuli were solid black circles presented on a white screen. The monitor was placed on the table with a movable track under it so that it could be moved to different distances. Right: the perceived size (measured via manual estimation) for each individual (indicated as each kind of symbols connected by gray lines) in Experiment 2. The black lines with symbols show the average across participants. **B**, Left: In Experiment 3, participants viewed the stimuli monocularly through a 1 mm pin-hole in complete dark. The stimuli were solid white circles presented on a black screen. Through the 1 mm hole, participants were only able to see part of the monitor but not the borders (blue dashed-line circle). Again, the monitor was moved to different distances. Right: the perceived size (measured via manual estimation) for each individual (shown as each gray line with symbols) in Experiment 3 during restricted viewing and their average results (black lines with symbols).

The display monitor was placed on a movable track mounted on a table. Viewing distance was manipulated by moving the display monitor to two different positions manually (**Figs. 2A left and 2B left**). To minimize the influence of any dynamic visual or oculomotor adjustments that would occur during the actual movement of the monitor on the visually evoked response induced by the test stimulus, the stimulus was not triggered by the experimenter until 1.5~2.5 s after the monitor had been moved and set in place at the far or near position. Thus, the long interval between the placement of the monitor and the onset of the stimulus ensured that all the distance cues were processed and any event-related visual and oculomotor signals evoked by the movement of the monitor had stabilized well before the stimulus was presented.

### Experiment 1

In this experiment, the stimulus was a black solid circle on a white background, and therefore its contrast and brightness changed minimally with viewing distance. Participants viewed the stimuli binocularly with the room lights on (full-viewing condition, **Fig. 2A** left). Because we manipulated the real distance of the stimulus display, many different cues to distance were available, including oculomotor adjustments (vergence, accommodation), pictorial cues, and binocular disparity, and were congruent with one another.

Participants were asked to identify whether the stimulus was the small one or the large one by pushing one of two keys. They all reported stimuli in both NS and FS as “small” (mean percentage of “small” response: NS, 99.11%; FS, 98.67%) and in both NL and FL as “large” (mean percentage of “small” response: NL, 98.28%; FL, 98.89%), suggesting that participants had size constancy in the full-viewing condition.

EEG signals were recorded from six electrodes (P3, P4, PZ, CP3, CP4 and CPZ) at the back of the head which typically yield the strongest visually evoked potentials (Chen et al., 2014; Chen, Yu, Zhu, Peng, & Fang, 2016). **Fig. 3A** shows the event-related potentials averaged across all six electrodes for each of the four conditions. The first visually evoked component C1, especially the initial portion of C1 between 56-70 ms after stimulus onset, is thought to be generated mainly by feedforward signals in V1 (Bao, Yang, Rios, He, & Engel, 2010; Clark, Fan, & Hillyard, 1994; Di Russo, Martínez, Sereno, Pitzalis, & Hillyard, 2002; Foxe & Simpson, 2002). Any feedback from higher-level visual areas will appear later in the event-related potentials (ERPs). The C1 component in the current experiment had a peak latency of 56 ms on average, which should have reflected the initial processing in V1 without trial-specific top-down influences being involved. If size constancy occurs at the initial stages of visual processing in V1, then stimuli of the same perceived size (i.e., stimuli with the same physical size but viewed at different distances) would evoke similar C1 amplitude. This was not the case, however. Instead, we found that only the NL stimulus, which had the largest retinal image size, evoked a significant C1 (t(1,15) = −3.86; p = 0.002), and the amplitude of C1 evoked by the NL stimulus was significantly larger than the one evoked by the FL stimulus, which had the same physical and perceived (but not retinal) size as the NL stimulus (t(1,16) = −3.08, p = 0.008), suggesting that C1 reflected the retinal image size, but not the physical size, of the stimulus.

**Fig 3.**
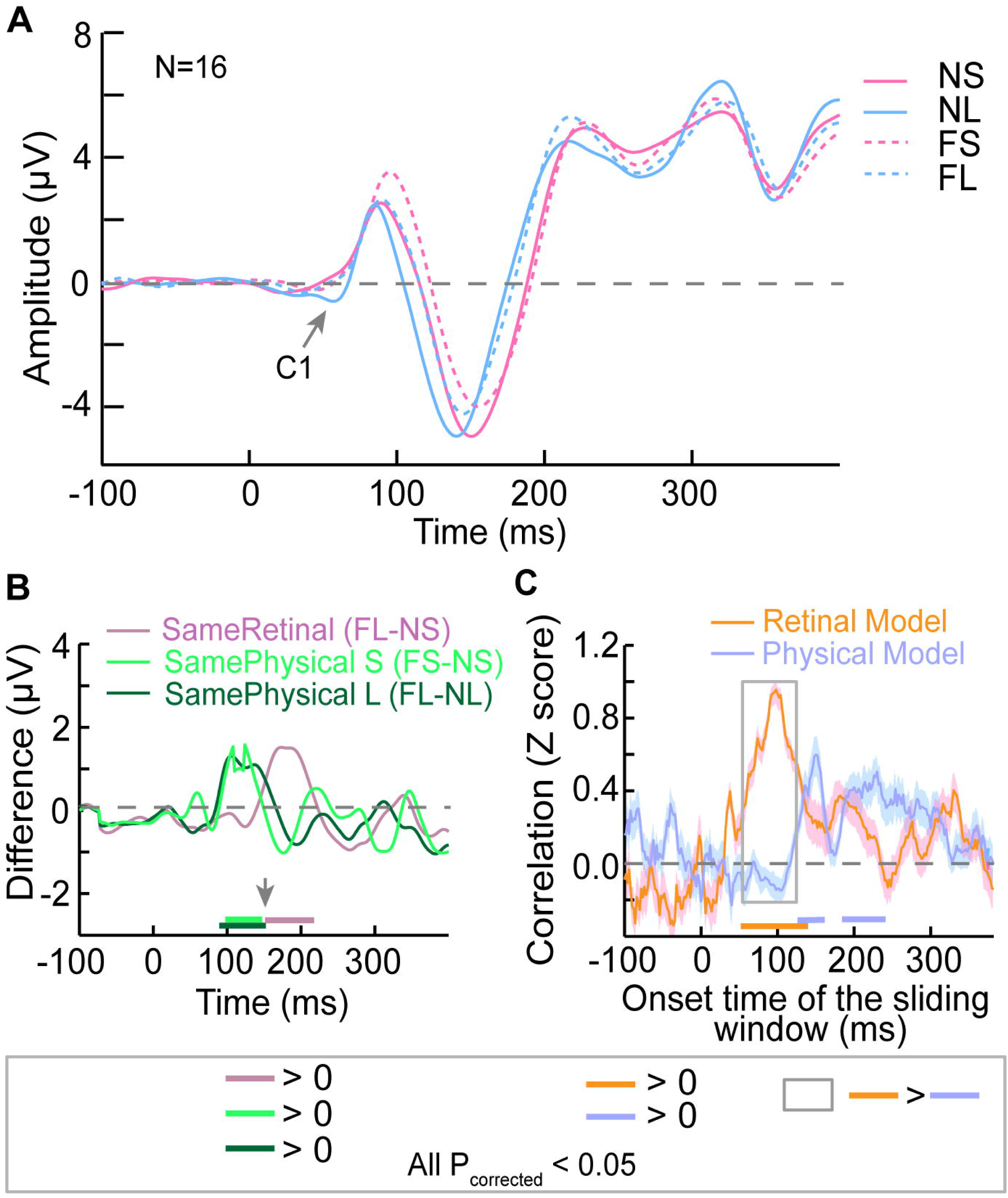
ERP results of Experiment 1 in which participants were asked to indicate whether the stimulus was large or small in the full-viewing condition. **A**, ERP curves that were first averaged across all six electrodes for each participant and then averaged across participants for each condition. **B,** The difference in amplitude between conditions that had the same retinal size (i.e., between NS and FL), and between conditions that had the same physical size (i.e., between FS and NS, and between FL and NL). The gray arrow points to approximately when the representation of retinal image size ended and when the signals began to change to represent the perceived size. **C**, The results of the representational similarity analysis (RSA). Each curve shows the time course of correlation between the similarity matrix of the neural model obtained from the ERP amplitude pattern and the similarity matrix of each of the size models (Retinal Size model and Physical Size model). The horizontal axis shows the start point of the 20-ms sliding time window. Shaded regions show standard error of the mean. The colored thick bars show when the values on each curve were significantly different from 0. The gray box shows when the two correlations were significantly different. All statistical point-by-point one sample t-tests or paired t-tests reported in this study were corrected using the cluster-based test statistic embedded in Fieldtrip toolbox (Monte Carlo method, p < 0.05).

As the ERP continued to unfold, the waveforms clustered in a way that reflected the physical size of the stimuli rather than their retinal image size [see **Fig. 3A**; note that the waveforms for the NL and the FL conditions (blue lines) overlap one another as do the waveforms for the NS and FS (pink lines)]. In short, the later components of the ERP appeared to show evidence of the operation of size constancy mechanisms.

To examine exactly when the transition from the representation of retinal image size to the representation of physical size occurred, we calculated the difference in the amplitude of the ERPs between conditions that had the same retinal image size (FL-NS) and conditions that had the same physical size (FS-NS and FL-NL). These difference scores, which are illustrated in **Fig. 3B**, revealed that waveforms for the stimuli with the same retinal image size (FL and NS) overlapped completely until 148 ms after stimulus onset at which point they began to separate, suggesting that before this time point the activity in visual cortex reflected only the size of the retinal image subtended by the stimulus [p_corrected_ < 0.05, corrected using a cluster-based test statistic (Monte Carlo) method embedded in Fieldtrip toolbox (Oostenveld, Fries, Maris, & Schoffelen, 2011); the same criterion was used for all time-course-related comparisons thereafter]. In contrast, the difference scores calculated for the ERPs in which the stimuli had the same physical size showed that the waveforms for the two small stimuli (FS and NS) began to overlap at 150 ms after stimulus onset and the waveforms for the two large stimuli (FL and NL) at 144 ms, suggesting that after these time points, the activity in visual cortex began to reflect the physical size of the visual stimuli. Taken together, these findings indicate that the activity in visual cortex reflected the retinal image size of visual stimuli until about 150 ms after stimulus onset but after that, began to represent the physical size of the stimuli.

The results reported above are all based on the amplitude difference averaged across all electrodes between each pair of conditions (i.e., pairs of conditions that had the same retinal image size or the same physical size). To further explore the temporal dynamics of processing associated with retinal image vs. physical size, we also performed a representational similarity analysis (RSA) based on the *patterns* of signals from all six electrodes within a 20-ms sliding time window. Each element of the similarity matrix for neural signals was the Pearson’s correlation between the EEG signal patterns of each pair of conditions (see Methods for details). If the visual signals were representing retinal image size, then the similarity matrix for the neural signals (neural model) should have a higher correlation with the similarity matrix for the retinal image size (retinal model, **Fig. 1B left**) than with the similarity matrix for the physical size (physical model, **Fig. 1B middle**). Consistent with our prediction, the RSA revealed that the neural model was significantly correlated with the retinal model before about 150 ms (**Fig. 3C**, see **Table 2** for details. Note: numbers in **Table 2** shows the *start* point of the 20-ms sliding window), and was significantly correlated with the physical model after about 124 ms. Importantly, the neural model was correlated more with the retinal model at 50~150 ms and correlatedmore with the physical model at a later window, although this difference did not survive correction for multiple comparisons (**Fig. 3C, Table 2**). Taken together, these results provide converging evidence that during the early stages of visual processing (within the first ~150 ms) the observed activity is locked to retinal image size but later on begins to reflect the real-world size of a visual stimulus.

**Table 1.** Time intervals (ms after stimulus onset) when the amplitude difference was significantly different from 0 with the p values corrected for multiple comparisons in brackets (cluster-based test statistic embedded in Fieldtrip toolbox (Monte Carlo method, p < 0.05).

|  | Same retinal size (FL-NS) | Same small physical size (FS-NS) | Same large physical size (FL-NL) |
| --- | --- | --- | --- |
| Exp 1 | 148-214 (0.006) | 88-150 (0.024) | 94-144 (0.002) |
| Exp 2 | 138-204 (0.002) | 90-162 (0.004) | 92-144 (0.01) |
| Exp 3 | 144-204 (0.04) | - | 154-350 (0.002) |

**Table 2.** The *start* time point of the sliding window (ms after stimulus onset) when the correlation (Fisher Z score) between the neural model and each of the size models was significantly different from 0 or the two correlations were significantly different from each other. The corrected p values (cluster-based test statistic embedded in Fieldtrip toolbox, Monte Carlo method, p < 0.05) is reported in brackets.

|  | Retinal<br>>0 | Physical<br>>0 | Perceived><br>0 | Retinal<br>> Physical | Physical<br>> Retinal | Retinal<br>> Perceived | Perceived<br>> Retinal |
| --- | --- | --- | --- | --- | --- | --- | --- |
| Exp 1 | 54-136<br>(0.002) | 124-158<br>(0.046)<br>182-238<br>(0.01) | - | 54-126<br>(0.002) | - | - | - |
| Exp 2 | 52-114<br>(0.002)<br>164-216<br>(0.012) | 180-246<br>(0.004) | 182-242<br>(0.002) | 66-124<br>(0.002) | - | 68-122<br>(0.002) | - |
| Exp 3 | 86-222<br>(0.002) | - | 82-114<br>(0.046)<br>128-156<br>(0.042)<br>222-382<br>(0.004) | 92-188<br>(0.002)<br>192-228<br>(0.016) | - | - | 236-268<br>(0.04) |

### Experiment 2

In Experiment 1, participants indicated whether the stimulus was large or small during EEG recording. One might argue that if they had not been asked to do a size-relevant task, the coding of physical size would emerge much later – or never. In other words, the post-150 ms overlap in the waveforms for the same physical size conditions might be due to nothing more than the fact that participants had only two choices in their behavioral response: small or large. To rule out these possibilities, we replicated the EEG protocol of Experiment 1, but asked participants to detect the onset of a non-stimulus visual target (an open circle) that was randomly interleaved with the experimental stimuli (solid circles) during the EEG recording. In addition, after the EEG recording, we also carried out a separate psychophysical test in which we asked participants to indicate the perceived size of each stimulus at each viewing distance by opening their thumb and index finger a matching amount (manual estimation task) (Chen, Jayawardena, & Goodale, 2015; Chen, Sperandio, & Goodale, 2015; Chen et al., 2018). The addition of the manual estimation task made it possible to measure any subtle differences in perceived size (i.e., size constancy) between participants, allowing us to calculate a perceived-size similarity matrix (**Fig. 1B right,** perceived model) for each participant and to then calculate the correlation between the perceived model and the neural model in a representational similarity analysis (RSA). Note that such an analysis was not possible in Experiment 1 because participants had been asked to categorize the stimuli as either large or small.

The manual estimation data confirmed that the participants on average showed size constancy (i.e. main effect of distance was not significant, F(1,13) = 0.002 p = 0.969; **Fig. 2A right**). Consistent with Experiment 1, only the NL condition, which had the largest retinal size, generated a significant C1 component (t (11) = −4.02, p = 0.002; **Fig. 4A**) and the C1 induced by the NL condition was significantly larger than that one induced by the FL condition (t (11) = 3.73, p = 0.003). The difference in amplitude between conditions that had the same retinal image size but different physical sizes (NS and FL) did not emerge until 138 ms after stimulus onset (**Fig. 4B**). The difference between conditions that had the same physical size (NL and FL) did not disappear until 144 ms for the large stimulus and 162 ms for the small stimulus (FS and FS) (**Fig. 4B**), which confirms the observation from **Fig. 4A** that after about 150 ms, the waveforms for conditions that had the same physical size started to overlap.

**Fig 4.**
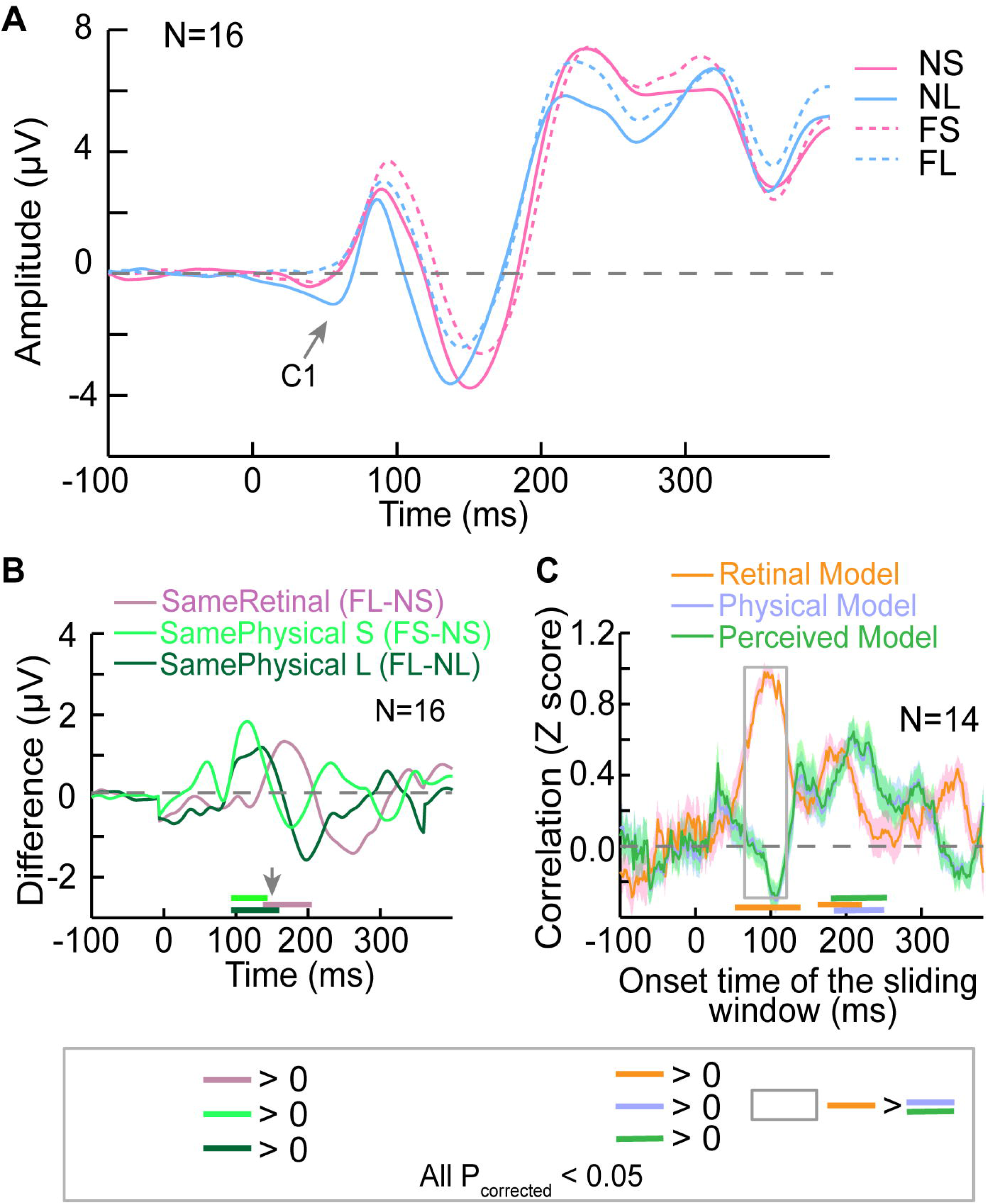
ERP results of Experiment 2 in which participants performed a size-irrelevant task (i.e., detect the onset of a non-testing stimulus) in the full-viewing condition. **A**, ERP curves that were first averaged across all six electrodes for each participant and then averaged across participants for each condition. **B**, The difference in ERP amplitude between conditions that had the same retinal size or the same physical size (same as **Fig 3B**). The gray arrow points to roughly when the size representation of retinal size ended and when the ERPs began to change to represent perceived size. **C**, The results of the RSA analysis. Each curve shows the time course of correlation between the similarity matrix of each size model and the similarity matrix of the neural model obtained from the ERP activation pattern. Shaded regions show standard error of the mean. The horizontal axis shows the start point of the 20-ms sliding time window. Again, the colored thick bars show when the difference was significantly different from 0. The gray box shows when the correlations between the Retinal Model and the Physical Model (and the Perceived Model) were significant.

The RSA also revealed a pattern of results that was similar to that seen in Experiment 1. First, the neural model was significantly correlated with the retinal model at an early stage, and was significantly correlated with the perceived and the physical model at a relatively later stage (see **Table 2** for the *start* time point of the sliding windows that showed significant difference between conditions; the correlation of the neural model with the physical-size model and the correlation of the neural model with the perceived-size model almost perfectly overlapped because almost all the participants showed size constancy, **Fig. 2A right**). Second and most importantly, the neural signals were correlated more with the retinal model than with the physical or the perceived size model before about 150 ms (**Table 2**, the start point of the 20-ms sliding window was from 66 ms to 124 ms after stimulus onset). All these results agree well with those in Experiment 1 and suggest that retinal image size, not perceived size, was encoded at the initial stage of visual processing and only later did the activity reflect the perception of stimulus size. The fact that the same timing was observed even when participants were performing a size-irrelevant task suggests that size-distance integration is to some extent automatic and independent of the task the participants were performing.

### Experiment 3

In the previous experiments, participants on average showed perfect size constancy in the full-viewing condition. We found strong and converging evidence that 150 ms after stimulus onset is the critical time point when the transition from coding retinal size to coding perceived size happens. In Experiment 3, we removed most of the cues to viewing distance, which we expected would disrupt size constancy and affect the perceived size of the stimulus. We then explored whether individual differences in the degree of disruption would be reflected in the grouping of the EEG components that unfolds after 150 ms.

Specifically, in Experiment 3, participants were asked to view the stimulus (a *white* solid circle on black screen, see Methods for more information) with their non-dominant eye through a 1- mm pinhole in an otherwise completely dark room (Chen et al., 2018; Holway & Boring, 1941) (i.e. restricted-viewing condition, **Fig. 2B left**), while performing a size-irrelevant detection task (the same task as in Experiment 2) during the EEG recording. In this case, no binocular distance cues were available. Pictorial cues were dramatically reduced as participants were able to see only a little bit of the background. In addition, the small pinhole prevented participants from using accommodation as a reliable cue to distance (Hennessy, Iida, Shiina, & Leibowitz, 1976). With such a restricted viewing condition, participants would have to rely mainly on retinal image size to judge object size; thus, a stimulus at the near distance would be perceived as larger than the same stimulus at the far distance because the stimulus would subtend a larger retinal image size at the near distance (Chen et al., 2018). This was confirmed by the manual estimates that participants provided in a separate behavioral test (without EEG recording) in the same pinhole viewing condition (**Fig. 2B right,** the main effect of distance, F = 91.344, p < 0.001). Nevertheless, it is important to point out that all the participants still knew whether the monitor was at the near or the far position, presumably on the basis of cues from the moving monitor when its position was changing – and from other cues, such as brightness and perhaps differences in the amount of background that was visible. As a result, the extent to which size constancy was disrupted would have depended on how well each individual could exploit the remaining distance cues. Indeed, there was considerable variability in size constancy across participants as shown in **Fig. 2B right**.

The peak of C1 in Experiment 3 occurred approximately 20 ms later than it did in experiments 1 and 2, probably because only one eye was being stimulated in this experiment (Mirzajani & Jafari, 2014). Nevertheless, consistent with Experiments 1 and 2, the NL stimulus, which had the largest retinal size, evoked the strongest C1 component (compared with the amplitude of the other three conditions, paired t-test, all t < 3.129, p < 0.006; **Fig. 5A**), again suggesting that retinal image size, not physical size, was driving the activity of the early ERP components. The waveforms for those conditions in which the stimulus subtended the same retinal image size (NS and FL) in Experiment 3 began to depart from each other around 144 ms after stimulus onset (**Fig. 5B**), just as they did in Experiments 1 and 2, but overall the waveforms did not show the same clear groupings according to physical size as they did in the two previous experiments. Instead, the waveform evoked by the NL stimulus began to separate from the FL stimulus approximately 154 ms after stimulus onset and never showed any overlap with FL, even though they had the same physical size. This pattern agrees with the fact that, under restricted viewing condition, the NL stimulus was perceived on average as being the largest stimulus of the four (**Fig. 2B, right**).

**Fig 5.**
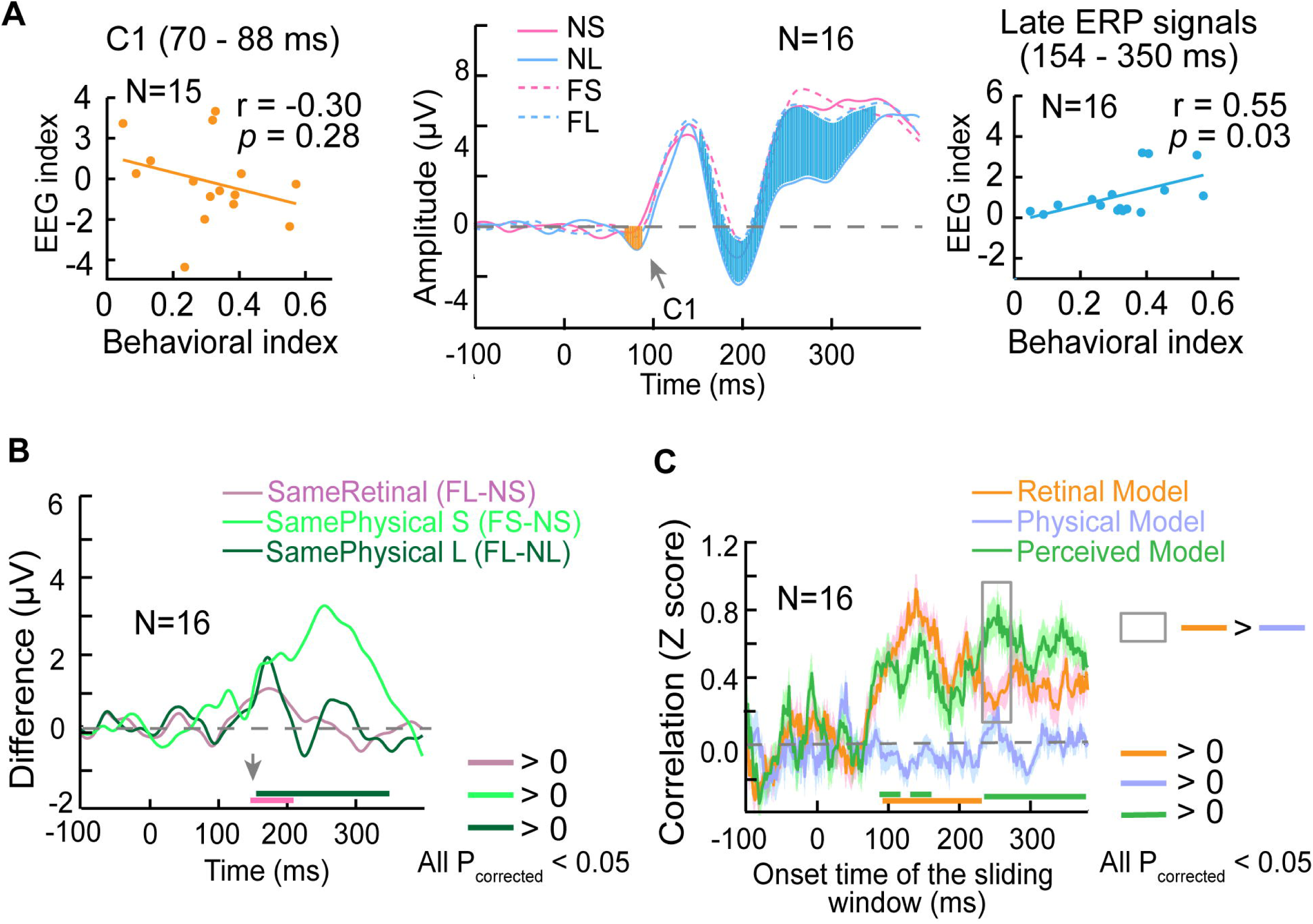
ERP results of Experiment 3 in which participants performed a size-irrelevant task (i.e. non-stimulus detection task) in the restricted-viewing condition. **A**, Middle: ERP curves that were first averaged across all six electrodes for each participant and then averaged across participants for each condition. Left: The scatter plot which shows the correlation between the amount of size-constancy disruption reflected in the behavioral performance (i.e. perceived size) and the amount of size-constancy disruption reflected in the earliest visual-evoked component C1 (i.e., the orange area in the middle figure). Right: The scatter plot showing the correlation between the amount of size-constancy disruption reflected in behavioral performance and the amount of size-constancy disruption reflected in the later ERP components (i.e., the blue area in the middle figure). **B**, The difference in ERP amplitude between conditions that had the same retinal size or the same physical size**. C,** RSA results. Each curve shows the time course of correlation between the similarity matrix of each size model and the similarity matrix of the neural model obtained from the ERP activation pattern. Shaded regions show standard error of the mean. Again, the colored thick bars in **B** and **C** show when the values on each curve were significantly different from 0 and the gray box show when the difference in the correlation of neural model with Retinal Model and with Perceived Model was significantly different.

Despite the evident disruption in size constancy on average across participants in the restricted-viewing paradigm, as was mentioned already, some participants did better than others in reporting the real size of the stimuli. Visual inspection revealed that, for participants whose size constancy was not disrupted or only slightly disrupted, the ERPs for the four conditions appeared to group according to the physical size as observed in both Experiments 1 and 2 (**Figs. 3A and 4A**). In contrast, for those participants whose size constancy was strongly disrupted, the waveform for the NL stimulus showed an increasingly large deviation from the waveform for the FL stimulus (and the other three conditions) after 150 ms. To quantify this, we calculated the correlation between behavioral reports and the waveforms of the ERP across participants. Specifically, we calculated a behavioral index (BI) of disruption in size constancy [(BI=ME_NL_-ME_FL_)/ ME_FL_, where ME indicates the manual estimate of perceived size]. We also calculated an EEG index (EI) of disruption in size constancy for the late component of the ERPs [(EI = (A_NL_-A_FL_)/ A_FL_, where A is the area separating the waveform and the x axis (i.e., amplitude = 0) where the waveforms for the stimuli in the NL and FL conditions are significantly different from one another (blue shaded area from 154 ms to 350 ms in **Fig. 5A, middle**)]. We found that there was indeed a significant correlation between BI and EI across participants (r = 0.55, p = 0.03; **Fig. 5A, right**). We also calculated a similar correlation between BI and EI for the early C1 component (the orange shaded area in **Fig. 5A, middle**) but the correlation was not significant (r = −0.30, p = 0.28; **Fig. 5A, left**), again suggesting that the variability in perceived size across participants is reflected in the later ERP components but not in C1.

Similar to Experiments 1 and 2, we also performed an RSA for Experiment 3. The correlation between the neural model and the physical model (**Fig. 5C)** was close to 0 throughout the whole post-stimulus interval, which is not surprising given that size constancy was disrupted to some degree for almost all the participants. In contrast, the retinal model and the perceived model were both highly correlated with the neural model from about 80 ms after stimulus onset (see **Table 2** for details). Although the perceived size was biased towards the retinal size in the restricted-viewing condition as shown in the behavioral data (**Fig. 2B right**), we found a trend in favor of the retinal model at the early stage (**Fig. 5C**, orange is above green) and a trend in favor of the perceived model at the later stage (**Fig. 5C**, green is above orange, see **Table 2** for statistical results). This again provides convincing evidence that retinal-size was being coded at the early stages of the ERP, whereas perceived size was represented at later stages.

## Discussion

The three experiments provide converging evidence that the computations underlying size constancy do not take place in the initial stages of visual processing in V1, or earlier. In other words, although the distance cues might modulate the spiking rate of a subset of neurons in LGN (Lal & Friedlander, 1990; Weyand & Malpeli, 1993), SC (Batini & Horcholle-Bossavit, 1979) and V1 (Rosenbluth & Allman, 2002; Trotter & Celebrini, 1999; Trotter et al., 1992; Weyand & Malpeli, 1993) at single-unit level, the integration with retinal image size still takes at least 150 ms to show perceived-size related activity at the neural population level (as revealed in our ERP results) in human participants.

It is important to note that unlike previous studies which used pictorial illusions (Fang et al., 2008; He et al., 2015; Liu et al., 2009; Murray et al., 2006; Ni et al., 2014) (e.g., the Ponzo illusion) projected on a screen at a fixed distance as stimuli, we changed the physical distance of the stimulus display from trial to trial, so that in the full-viewing condition in Experiments 1 and 2 there was a large range of distance cues, including oculomotor, binocular, and monocular cues, which were entirely congruent with one another. More importantly, the long interval after the monitor had been set in place provided enough time for the distance cues to be well processed before the stimulus onset, so that the distance information could theoretically be integrated with retinal information about the test stimulus as soon as the stimulus was presented. For all these reasons, the time we identified as the transition point from the coding of retinal image size to the coding of perceived size, which occurred at approximately 150 ms after stimulus onset, is probably the earliest possible time point at which the integration of retinal image size and viewing distance information can take place. Interestingly, the same time interval was required to compute perceived size in Experiment 3 when visual cues to distance were degraded (but still congruent) and participants showed large individual differences in size constancy judgments. Taken together, these results suggest that 150 ms is an interval that may be required for the integration of distance information with retinal image size no matter what (congruent) visual or oculomotor cues are available.

Two other studies have also examined the timing of activity in visual areas related to the computation of perceived size. Ni et al. (2014) found that the position of receptive fields of neurons in V1 in the monkey were influenced by the position of a visual stimulus on a Ponzo illusion background – and that this effect was evident extremely early (about 30 ms after stimulus onset). Of course, conductance times in the monkey brain will always be much shorter than those in the human brain because of the large difference in brain size. Moreover, the modulation of activity that was observed in a subset of neurons using single-unit recording does not necessarily represent the population level coding of perceived size in V1. Indeed, an ERP study by Liu et al. (2009), which also used the same illusory display in human participants, found that the modulation of the signal occurred much later, around 240-260 ms, a time that is even later than the 150 ms we observed here. This difference may be related to the fact that all the distance cues in our study were congruent, whereas in Liu et al.’s study the pictorial cues were in conflict with the vergence, accommodation, and binocular disparity cues, perhaps leading to a delay in the integration of the pictorial cues with retinal image size. In other words, by eliminating the conflict between distance cues, we revealed the real timing of the transition from the coding of retinal image size to coding of perceived size.

Because the 150 ms required for the size-distance integration is consistent with the time that is typically required (80 to 150 ms after stimulus onset) for the recurrent feedback from higher-order visual areas to V1 (Wyatte, Jilk, & O'Reilly, 2014), it is possible that the representation of perceived size in V1 observed in previous fMRI studies (Fang et al., 2008; He et al., 2015; Murray et al., 2006; Sperandio et al., 2012) also reflected recurrent processing. Recurrent feedback to V1 has been shown to be critical for feature binding (Bouvier & Treisman, 2010; Koivisto & Silvanto, 2011). In a similar fashion, such feedback could be used to integrate distance information with retinal image size to calculate the real-world size of objects, and subsequently, integrate real-world size with other object features, such as shape, colour, and visual texture. Indeed, it is worth noting that accounts of feature integration have almost entirely ignored object size, perhaps because only images presented on a display at a fixed distance rather than real objects presented at different distances have been employed in these studies.

Interestingly, size constancy is not only observed in perceptual judgments, but is also observed in grasping movements – that is, within a comfortable reaching space, people typically use the same grip aperture to grasp an object regardless of viewing distance. Here, we found that size constancy does not happen in the initial stage of V1 (or even earlier) even when real distance was changed. This finding agrees with the observation that proprioceptive distance cues make a larger contribution to size constancy in grasping than to size constancy in perception when visual cues are limited (Chen et al., 2018), which also suggests that size-distance integration does not happen early in V1 or even before, but may happen in the dorsal visual stream and the motor/premotor cortex for grasping and in the ventral visual cortex for perception. Moreover, it has been suggested that efference copy information from vergence (and theoretically accommodation) is conveyed from the superior colliculus (via thalamic nuclei) to the frontal eye fields and to visuomotor areas in the posterior parietal cortex, completely by-passing the geniculostriate pathway altogether (Sommer & Wurtz, 2008). Therefore, it is likely that although the integration of retinal image size and distance information takes at least 150 ms for perception, some distance information could be conveyed to visuomotor networks in the dorsal stream quickly (Chen et al., 2007; Foxe & Simpson, 2002) to mediate size constancy for grasping. Additional support for this idea comes from studies showing that patients with lesions of V1 can scale the opening of their grasping hand to the size and orientation of goal objects (Carey, Dijkerman, & Milner, 1998; Carey, Harvey, & Milner, 1996; Prentiss, Schneider, Williams, Sahin, & Mahon, 2018; Whitwell, Striemer, Nicolle, & Goodale, 2011), even though they do not perceive those objects.

In sum, our study provides accurate temporal information about how representation of size changes with real distance in natural circumstances. The finding clarifies the role of V1 in size constancy and has implications in any cognitive processing and motor controls that involve size-distance integration.

## Materials and Methods

### Participants

Seventeen participants (7 males, 10 females) took part in Experiment 1. One participant’s (male) data was discarded because of strong noise in his EEG signals. Sixteen participants (5 males, 11 females) took part in the EEG portion of Experiment 2, but only 14 of them took part in the behavioral portion of the experiment. Two participants were unable to complete the behavioral portion due to lack of time in the testing session. Sixteen participants (6 males, 10 females) took part in both the EEG and the behavioral size estimation portions of Experiment 3. All were right handed and had no history of neurological impairments. Most of participants aged between 18 and 30 years old except for two participants in Experiment 3, who were 45 and 52 years old, respectively. Participants in Experiments 1 and 2 had either normal or corrected-to-normal visual acuity. All participants in Experiment 3 had normal visual acuity. Informed consent was obtained from all subjects according to procedures and protocols approved by the Health Sciences Research Ethics Board at The University of Western Ontario.

### Stimuli and setup

In Experiments 1 and 2, the stimuli were black solid circles with a diameter of 4 cm (i.e. ‘Small’ or ‘S’) or 8 cm (i.e. ‘Large’ or ‘L’). They were presented in the center of a screen with a white background (**Fig 2A**). The screen was mounted on a movable track so that the experimenter could move it to a near (28.5 cm, ‘N’) or a far viewing distance (57 cm, ‘F’). In these two experiments, the near-small (NS) and far-large (FL) stimuli had the same retinal size; the near-small (‘NS’) and far-small (‘FS’) stimuli had the same physical size, so did the near-large (‘NL’) and far-large (‘FL’) stimuli. We used black circles on a white background, instead of white circles on a black background as stimuli, so that the brightness and perceived contrast would not vary with the viewing distance. We used solid circles, instead of gratings or other complex objects as stimuli, to avoid any confound of differences in spatial frequency at different viewing distances. There was a fixation point (a red dot) on the center of the screen throughout the experiments. Participants were seated in front of the screen with their chin on a chinrest. These two experiments were performed with the room lights on and under binocular viewing conditions (i.e., full-viewing condition).

In Experiment 3, the same design (2 sizes × 2 distances) was adopted. The room was completely dark and participants looked at the stimuli through a 1 mm hole on the pin-hole glasses with their non-dominant eye (i.e., restricted-viewing condition). Unlike Experiments 1 and 2, in which black solid circles were presented on a white screen, in Experiment 3, the stimuli were *white* solid circles presented on a *black* background. The reason for introducing this change was that if we used black circles as stimuli and white screen as background in Experiment 3, participant would be able to see the boundary of the circular field of view clearly when they wore pin-hole glasses. The relative size between the circular stimuli and the area they could see through the pin-hole would have provided them with information regarding the size of the stimuli, which would have made it impossible to disrupt size constancy. Because white circles, instead of black circles, were used as stimuli to make sure that size constancy could be disrupted in the restricted-viewing condition (see Method for details), the brightness and contrast of the stimulus would have varied with viewing distance, which might explain why the waveforms of conditions were not as well organized as those in Experiments 1 and 2 at the second ERP component (peak at about 150 ms after stimulus onset). When a white stimulus was presented on a black screen and was viewed through a 1 mm hole with one eye in darkness, all the binocular cues and most of the contextual cues were not available anymore. They could not see the frame of the monitor. The blur was removed and accommodation could not provide valid distance information (Hennessy et al., 1976). Previous studies have shown that size constancy could be disrupted effectively with these stimuli and setup (Holway & Boring, 1941). Nevertheless, because participants could still indicate the distance based on the movement of the monitor, the extent to which size constancy was disrupted would depend on how well each individual could use the remaining distance cues.

### Procedure

In Experiment 1, participants were asked to indicate whether a solid circle was small or large regardless of distance by pressing two keys (“1” for small and “2” for large) during EEG recording. At the beginning of each trial, the experimenter was cued with a letter, either ‘N’ or ‘F’, that appeared at the corner of the screen to indicate whether the viewing distance of a specific trial would be near or far (note: the participants could not see the letter). The experimenter who sat beside the monitor would move the monitor to the near or far position, accordingly. 1.5 ~2.5 s after the screen was moved to the right position, the experimenter pushed a key to trigger the presentation of the stimulus. The stimulus was presented on the screen for 0.2 s. Participants were asked to maintain fixation at the fixation point throughout the experiment. There were 100 trials in each run, with 25 trials for each condition.

In Experiment 2, the protocol of the EEG trials was the same as that described for Experiment 1 with two exceptions. First, during EEG recording in each run, there were 10 additional trials in which the stimulus was an open circle, rather than a solid circle. Participants were asked to push a key (“0”) as soon as they saw the open circle (i.e., target-detection task). Second, in addition to the EEG trials, 14 out of the 16 participants also performed a behavioral task in which they were asked to open their thumb and index finger to indicate the *perceived* size of the stimulus (manual estimation task) (Chen, Jayawardena, et al., 2015; Chen, Sperandio, et al., 2015; J. Chen et al., 2018). The distance between the finger and thumb was then measured with a measuring tape. This psychophysical measure was taken after the EEG session. Participants completed 4-5 psychophysical blocks depending on the time available, with 2 manual estimates for each of the four conditions in each block.

In Experiment 3, the same EEG protocol was used as reported above. Participants also performed a detection task during EEG recording and also performed a separate behavioral testing session. As mentioned above, the key difference between this experiment and Experiment 2 was that the stimulus was a white solid circle on a black background and participants viewed the stimulus monocularly with their non-dominant eye through a 1 mm hole in the dark (i.e., restricted-viewing condition). In addition, unlike Experiment 2, the psychophysical blocks were performed before any EEG recordings and after every four EEG runs, in case the perceptual experience of size changed over EEG runs.

In all experiments, the order of the four conditions was randomized on a trial-by-trial basis. Participants completed between 8 and 14 runs of EEG recording depending on the time available, for a total of 200-300 repetitions for each condition. Each experiment lasted between 3 and 4 hours.

It should be noted that size constancy was not affected by the restricted-viewing condition to the same extent across participants, probably because of individual differences in their ability to use residual depth cues (e.g. vibration or auditory cues provided by the movement of the monitor, or changes in the brightness of the white stimulus) to enable size constancy. [In another study from our lab in which we moved a sphere, rather than a monitor to different location on a table, we were able to successfully disrupt size constancy in all participants using the same restricted-viewing condition (Chen et al., 2018)]. We noticed this issue after we completed the EEG recording and behavioral testing of the first participant. Because the purpose of this investigation was to explore the neural correlates of perceived size when size constancy was disrupted, we performed additional psychophysical tests to exclude those participants whose size constancy was not affected at all by the restricted viewing conditions. Thirty-one participants took part in these additional tests in which they were required to manually estimate the size of the circle under the restricted-view condition. The size constancy of 15 out of the 31 participants originally tested was affected to some extent, and therefore only these 15 participants were included in Experiment 3 together with the first tested participant.

### EEG measurements

Scalp EEG was collected using NeuroScan Acquire 4.3 recording system (Compumedics) from 32 Ag/AgCl electrodes positioned according to the extended international 10 – 20 EEG system. Vertical electro-oculogram (VEOG) was recorded from two electrodes placed above or below the left eye. Horizontal EOG (HEOG) was recorded from two electrodes placed at the outer canthus of the left and the right eyes. Because we were interested in the six electrodes at the parietal and occipital part of the scalp (i.e., CP3, CPZ, CP4, P3, PZ, and P4) that have been reported to reflect visual processing (Luck, 2005), we always kept the impedance of these six electrodes below 10 kΩ. We also tried to keep the impedance of the other electrodes as low as possible, but this revealed to be impossible for all participants due to the long duration of the EEG session (> 3 hours). EEG was amplified with a gain of 500 K, band pass filtered at 0.05 – 100 Hz, and digitized at a sampling rate of 500 Hz. The signals on these electrodes were referenced online to the electrode on the nose.

## Data Analysis

### ERP data Preprocessing

Offline data analysis was performed with NeuroScan Edit 4.3 (Compumedics) and MATLAB R2014 (Mathwork). The EEG data was first low-pass filtered at 30 Hz, and then epoched starting at 100 ms before the stimulus onset and ending 400 ms after stimulus onset. Each epoch was baseline-corrected against the mean voltage of the 100 ms pre-stimulus interval. The epochs contaminated by eye blinks, eye movements, or muscle potentials exceeding ± 50 μV at any electrode were excluded from the average.

### Amplitude and latency analyses of ERP components

For the event-related potential (ERP) analysis, the remaining epochs after artifact rejection were averaged for each condition. Preliminary analyses revealed that the activity pattern of the four conditions in all 6 electrodes (i.e., CP3, CPZ, CP4, P3, PZ, and P4) were similar. Therefore, only the ERP amplitude and latency results that were averaged across these six electrodes were reported. The peak amplitude and latency of each component were acquired for each condition and each participant.

### Representational similarity analysis (RSA)

To examine at what time the brain activity was representing the retinal size, physical size or perceived size, we calculated the correlation between the similarity matrix revealed in neural signals (i.e., ERP amplitude) and similarity matrices for the retinal size, physical size and the perceived size, respectively, for each sliding window (10 time points, i.e., 20 ms) with the first point of the window moving from −100 ms to 382 ms. The element of the similarity matrix for the neural model (i.e., EEG signals) was set as the Fisher-Z correlation coefficient between the EEG patterns for each pair of conditions at a specific time window. Each EEG patterns included 60 elements (10 time points × 6 electrodes).

The similarity matrices for the retinal size and the physical size are shown in **Fig. 1B**. The similarity between two conditions was set as 1 if the retinal size or the physical size was the same, but was set as 0 if the retinal size or the physical size was different. These matrices were fixed across participants. The similarity matrix for perceived size was calculated for each individual (see **Fig 1B** for an example). Each element of the matrix was obtained by first calculating the perceived size difference between two conditions, and then multiplying the obtained value by −1. For Experiment 1, no perceived size data was collected for each individual, and therefore only retinal-size model and physical size model were tested.

To obtain an unbiased measurement of the correlation between the neural model and the size model, we used a procedure similar to the n-folded cross-validation that was commonly used in pattern recognition analysis. Specifically, we first randomly sampled half group of trials from the whole set of ERP trials for each condition, then we averaged the ERPs of the sampled trials. The averaged ERPs were used to calculate the correlation coefficients between the EEG patterns of each pair of conditions (i.e., the elements of the neural model) at a specific time window and to calculate the correlation between the obtained neural model and size model. This procedure was repeated 50 times. The 50 correlation coefficients between the neural model and size model were first converted to Fisher-Z scores, and were then averaged to arrive at the reported correlation results.

### Correlation between size constancy disruption index calculated in perceptual judgments and in ERP components

In Experiment 3, to test which ERP component reflected the individual variability in size-constancy disruption, we calculated the correlation between the amounts of size-constancy disruption measured behaviourally and the amount of size-constancy disruption measured in the ERP components across individuals.

The behavioral size-constancy disruption index (BI) was defined as
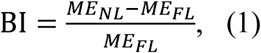

where ME indicates manual estimate.

To specifically examine whether the disruption of size constancy was reflected in the early visual component C1 or the late ERP component, the size constancy disruption in ERP was calculated separately for C1 and the late ERP component. The EEG size constancy disruption index (EI) was defined as
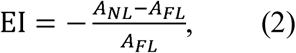

where “A” stands for the area under the curve (i.e., between the curve and the x axis) in a specific interval. For C1, this interval was when the C1 amplitudes in the NL condition were significantly higher than the 25% of the peak amplitude of the C1 in the same condition. In the current case, the interval was between 78-90 ms after stimulus onset. For the late EEG component, the interval was when the amplitude of NL was significantly different from the FL condition (154-350 ms). The large size, but not the small size, was used to calculate the behavioral and EEG size-constancy disruption indices because the size constancy disruption (i.e., the difference in perceived size or in ERP amplitude between near and far distances) was more evident and reliable in the large size condition than in the small size condition in both behavioral and EEG results. Pearson correlation was calculated to test whether or not the correlation between behavioral performance and neural signals were significant. For C1, one outlier (beyond +/-5 SD) was excluded.

### Statistical Analysis

To examine whether or not there was size constancy, repeated ANOVAs with size and distance as main factors were carried out to reveal whether or not the main effect of distance was significant. To compare the amplitude of C1 component evoked by different conditions, paired t-tests were performed on the peak value of the C1 amplitude. To search intervals when there were significant differences between each time course and 0 or between two time courses, paired t-tests were conducted point-by-point, and then were corrected for multiple comparisons using the cluster-based test statistic embedded in Fieldtrip toolbox (Monte Carlo method, p < 0.05). For the RSA results and the correlation between BI and EI results, all statistical comparison were conducted on the Fisher Z scores of the Pearson correlation coefficients.

## Acknowledgements

We are grateful to Amratha Chandrakumar and Jason Kim for their help with data collection. This research was supported by a Discovery Grant from the Natural Sciences and Engineering Research Council of Canada (MAG) and a Canadian Institute for Advanced Research Grant (MAG).

## Notes

**Competing financial interests** The authors declare no competing financial interests.

## References

Bao, M., Yang, L., Rios, C., He, B., & Engel, S. A. (2010). Perceptual Learning Increases the Strength of the Earliest Signals in Visual Cortex. Journal of Neuroscience, 30(45), 15080-15084.

Batini, C., & Horcholle-Bossavit, G. (1979). Extraocular muscle afferents and visual input interactions in the superior colliculus of the cat. In Progress in Brain Research (Vol. 50, pp. 335-344): Elsevier.

Blakemore, C., Garner, E. T., & Sweet, J. A. (1972). The site of size constancy. Perception, 1(1), 111-119. doi:10.1068/p010111

Bouvier, S., & Treisman, A. (2010). Visual Feature Binding Requires Reentry. Psychological Science, 21(2), 200-204. doi:10.1177/0956797609357858

Carey, D. P., Dijkerman, H. C., & Milner, A. D. (1998). Perception and action in depth. Consciousness and cognition, 7(3), 438-453.

Carey, D. P., Harvey, M., & Milner, A. D. (1996). Visuomotor sensitivity for shape and orientation in a patient with visual form agnosia. Neuropsychologia, 34(5), 329-337. doi:http://dx.doi.org/10.1016/0028-3932(95)00169-7

Chen, C.-M., Lakatos, P., Shah, A. S., Mehta, A. D., Givre, S. J., Javitt, D. C., & Schroeder, C. E. (2007). Functional Anatomy and Interaction of Fast and Slow Visual Pathways in Macaque Monkeys. Cerebral Cortex, 17(7), 1561-1569. doi:10.1093/cercor/bhl067

Chen, J., He, Y., Zhu, Z., Zhou, T., Peng, Y., Zhang, X., & Fang, F. (2014). Attention-Dependent Early Cortical Suppression Contributes to Crowding. The Journal of Neuroscience, 34(32), 10465-10474. doi:10.1523/jneurosci.1140-14.2014

Chen, J., Jayawardena, S., & Goodale, M. A. (2015). The effects of shape crowding on grasping. Journal of Vision, 15(3), 1-9. doi:10.1167/15.3.6

Chen, J., Sperandio, I., & Goodale, M. A. (2015). Differences in the effects of crowding on size perception and grip scaling in densely cluttered 3-D scenes. Psychological Science, 26(1), 58-69. doi:10.1177/0956797614556776

Chen, J., Sperandio, I., & Goodale, M. A. (2018). Proprioceptive Distance Cues Restore Perfect Size Constancy in Grasping, but Not Perception, When Vision Is Limited. Current Biology, 28, 1-6. doi:10.1016/j.cub.2018.01.076

Chen, J., Yu, Q., Zhu, Z., Peng, Y., & Fang, F. (2016). Spatial summation revealed in the earliest visual evoked component C1 and the effect of attention on its linearity. Journal of Neurophysiology, 115(1), 500-509. doi:10.1152/jn.00044.2015

Clark, V. P., Fan, S., & Hillyard, S. A. (1994). Identification of early visual evoked potential generators by retinotopic and topographic analyses. Human Brain Mapping, 2(3), 170-187.

Di Russo, F., Martínez, A., Sereno, M. I., Pitzalis, S., & Hillyard, S. A. (2002). Cortical sources of the early components of the visual evoked potential. Human Brain Mapping, 15(2), 95-111.

Dobbins, A. C., Jeo, R. M., Fiser, J., & Allman, J. M. (1998). Distance Modulation of Neural Activity in the Visual Cortex. Science, 281(5376), 552-555. doi:10.1126/science.281.5376.552

Fang, F., Boyaci, H., Kersten, D., & Murray, S. O. (2008). Attention-Dependent Representation of a Size Illusion in Human V1. Current Biology, 18(21), 1707-1712.

Foxe, J. J., & Simpson, G. V. (2002). Flow of activation from V1 to frontal cortex in humans. Experimental Brain Research, 142(1), 139-150.

He, D., Mo, C., Wang, Y., & Fang, F. (2015). Position shifts of fMRI-based population receptive fields in human visual cortex induced by Ponzo illusion. Experimental Brain Research, 233(12), 3535-3541. doi:10.1007/s00221-015-4425-3

Hennessy, R. T., Iida, T., Shiina, K., & Leibowitz, H. (1976). The effect of pupil size on accommodation. Vision Research, 16(6), 587-589.

Holway, A. H., & Boring, E. G. (1941). Determinants of apparent visual size with distance variant. The American Journal of Psychology, 21-37.

Humphrey, N. K., & Weiskrantz, L. (1969). Size constancy in monkeys with inferotemporal lesions. Quarterly Journal of Experimental Psychology, 21(3), 225-238. doi:10.1080/14640746908400217

Koivisto, M., & Silvanto, J. (2011). Relationship between visual binding, reentry and awareness. Consciousness and cognition, 20(4), 1293-1303. doi:https://doi.org/10.1016/j.concog.2011.02.008

Lal, R., & Friedlander, M. J. (1990). Effect of passive eye position changes on retinogeniculate transmission in the cat. Journal of Neurophysiology, 63(3), 502-522.

Liu, Q., Wu, Y., Yang, Q., Campos, J. L., Zhang, Q., & Sun, H. J. (2009). Neural correlates of size illusions: an event-related potential study. NeuroReport, 20(8), 809-814. doi:10.1097/WNR.0b013e32832be7c0

Luck, S. J. (2005). An Introduction to the Event-Related Potential Technique. Cambridge, MA: Massachusetts Institute of Technology.

Marg, E., & Adams, J. (1970). Evidence for a neurological zoom system in vision from angular changes in some receptive fields of single neurons with changes in fixation distance in the human visual cortex. Cellular and Molecular Life Sciences, 26(3), 270-271.

Masson, G. S., Busettini, C., & Miles, F. A. (1997). Vergence eye movements in response to binocular disparity without depth perception. Nature, 389(6648), 283-286.

Mirzajani, A., & Jafari, A. (2014). The effect of binocular summation on time domain transient VEP wave’s components. Razi Journal of Medical Sciences, 21(123), 29-35.

Murray, S. O., Boyaci, H., & Kersten, D. (2006). The representation of perceived angular size in human primary visual cortex. Nature Neuroscience, 9(3), 429-434.

Ni, Amy M., Murray, Scott O., & Horwitz, Gregory D. (2014). Object-Centered Shifts of Receptive Field Positions in Monkey Primary Visual Cortex. Current Biology, 24(14), 1653-1658. doi:http://dx.doi.org/10.1016/j.cub.2014.06.003

Oostenveld, R., Fries, P., Maris, E., & Schoffelen, J.-M. (2011). FieldTrip: Open Source Software for Advanced Analysis of MEG, EEG, and Invasive Electrophysiological Data. Computational Intelligence and Neuroscience, 2011, 9. doi:10.1155/2011/156869

Pooresmaeili, A., Arrighi, R., Biagi, L., & Morrone, M. C. (2013). Blood oxygen level-dependent activation of the primary visual cortex predicts size adaptation illusion. The Journal of Neuroscience, 33(40), 15999-16008. doi:10.1523/JNEUROSCI.1770-13.2013

Prentiss, E. K., Schneider, C. L., Williams, Z. R., Sahin, B., & Mahon, B. Z. (2018). Spontaneous in-flight accommodation of hand orientation to unseen grasp targets: A case of action blindsight. Cognitive Neuropsychology, 1-9. doi:10.1080/02643294.2018.1432584

Richards, W. (1968). Spatial remapping in the primate visual system. Kybernetik, 4(4), 146-156.

Richards, W. (1971). Size-distance transformations. In Zeichenerkennung durch biologische und technische Systeme/Pattern Recognition in Biological and Technical Systems (pp. 276-287): Springer.

Rosenbluth, D., & Allman, J. M. (2002). The Effect of Gaze Angle and Fixation Distance on the Responses of Neurons in V1, V2, and V4. Neuron, 33(1), 143-149. doi:http://dx.doi.org/10.1016/S0896-6273(01)00559-1

Schwarzkopf, D. S., Song, C., & Rees, G. (2011). The surface area of human V1 predicts the subjective experience of object size. Nature Neuroscience, 14(1), 28-30.

Smith, J., & Marg, E. (1975). Zoom neurons in visual cortex: Receptive field enlargements with near fixation in monkeys. Experientia, 31(3), 323-326.

Sommer, M. A., & Wurtz, R. H. (2008). Brain circuits for the internal monitoring of movements. Annu. Rev. Neurosci., 31, 317-338.

Sperandio, I., & Chouinard, P. A. (2015). The mechanisms of size constancy. Multisensory research, 28(3-4), 253-283.

Sperandio, I., Chouinard, P. A., & Goodale, M. A. (2012). Retinotopic activity in V1 reflects the perceived and not the retinal size of an afterimage. Nat Neurosci, 15(4), 540-542. doi:10.1038/nn.3069

Trotter, Y., & Celebrini, S. (1999). Gaze direction controls response gain in primary visual-cortex neurons. Nature, 398(6724), 239-242.

Trotter, Y., Celebrini, S., Stricanne, B., Thorpe, S., & Imbert, M. (1992). Modulation of neural stereoscopic processing in primate area V1 by the viewing distance. Science, 257(5074), 1279-1281.

Ungerleider, L., Ganz, L., & Pribram, K. (1977). Size constancy in rhesus monkeys: effects of pulvinar, prestriate, and inferotemporal lesions. Experimental Brain Research, 27(3-4), 251-269.

Weyand, T. G., & Malpeli, J. G. (1993). Responses of neurons in primary visual cortex are modulated by eye position. Journal of Neurophysiology, 69(6), 2258-2260. doi :10.1152/jn.1993.69.6.2258

Whitwell, R. L., Striemer, C. L., Nicolle, D. A., & Goodale, M. A. (2011). Grasping the non-conscious: Preserved grip scaling to unseen objects for immediate but not delayed grasping following a unilateral lesion to primary visual cortex. Vision Research, 51(8), 908-924. doi:http://dx.doi.org/10.1016/j.visres.2011.02.005

Wyatte, D., Jilk, D. J., & O'Reilly, R. C. (2014). Early recurrent feedback facilitates visual object recognition under challenging conditions. Frontiers in Psychology, 5, 674.

Wyke, M. (1960). Alterations of size constancy associated with brain lesions in man. Journal of neurology, neurosurgery, and psychiatry, 23(3), 253.

